# Geometry of visual working memory information in human gaze patterns

**DOI:** 10.1101/2022.11.17.516917

**Authors:** Juan Linde-Domingo, Bernhard Spitzer

**Author notes:** Correspondence to: Juan Linde-Domingo or Bernhard Spitzer.

## Abstract

Stimulus-dependent eye movements have been recognized as a potential confound in decoding visual working memory information from neural signals. Here, we combined eye-tracking with representational geometry analyses to uncover the very information in miniature gaze patterns while participants (n = 41) were cued to maintain visual object orientations. Although participants were discouraged from breaking fixation via real-time feedback, small gaze shifts (< 1 degree) robustly encoded the to-be-maintained stimulus orientation, with evidence for encoding two sequentially presented orientations at the same time. While the orientation encoding upon stimulus presentation was object-specific, it changed to a more object-independent format during cued maintenance, particularly when attention had been temporarily withdrawn from the memorandum. Finally, categorical reporting biases increased after unattended storage, with indications of biased gaze geometries emerging already during the maintenance periods prior to behavioral reporting. These findings disclose a wealth of information in gaze patterns during visuospatial working memory, and suggest systematic changes in representational format when memory contents have been unattended.

## Introduction

Working Memory (WM) enables observers to actively keep stimulus information “on the mind” for upcoming tasks. A key question in understanding WM function is which aspects of a task-relevant stimulus it retains, and in which format(s). Upon removal of a stimulus from sight, sensory systems briefly retain a detailed sensory memory of the just-removed (e.g., visual) input. Sans active maintenance, these rich “photographic” memories decay rapidly within a few hundreds of milliseconds (Sperling, 1960; but see e.g. Brady et al., 2008). It is widely assumed that only a limited amount of information can be accurately maintained in WM (Bays et al., 2009; Cowan, 2001; Luck & Vogel, 1997; Ma et al., 2014). However, despite intense research, the very nature of the information that WM maintains remains poorly understood.

In neuroscientific experiments examining the representation of visual WM information in the brain, often only a single stimulus feature needs to be reported after a delay (e.g., the orientation of a visual grating; Christophel et al., 2018; Harrison & Tong, 2009; Rademaker et al., 2019; Serences et al., 2009). At one extreme, such tasks could be solved by sustaining a concrete visual memory of the stimulus and its visual details. Various human neuroimaging studies have shown that WM contents can be decoded from early visual cortices (Harrison & Tong, 2009; Riggall & Postle, 2012; Serences et al., 2009), which seems consistent with storage in a sensory format (but see Kwak & Curtis, 2022). At the other extreme, many WM tasks can also be solved with a high-level abstraction of the task-relevant stimulus parameter only, such as its orientation, speed, or color (e.g., Kwak & Curtis, 2022; Romo et al., 1999; Spitzer & Blankenburg, 2012; Vergara et al., 2016). Such abstractions may also be recoded into pre-existing categories (such as “left”, “slow”, or “green”; e,g. Bae et al., 2015; Hardman et al., 2017), which may result in memory reports that are biased (see also Taylor & Bays, 2018) but still sufficient to achieve one’s behavioral goals. Abstraction may render working memories more robust, afford transfer across tasks, and may massively reduce the amount of information that needs to be maintained (see also Hardman et al., 2017; Ricker & Cowan, 2010; Vergauwe et al., 2014).

However, progress in understanding the temporal dynamics of WM abstraction has thus far been limited. Few studies have examined the extent to which neural WM representations generalize (or not) across different stimulus inputs (Christophel, Allefeld, et al., 2018; Kwak & Curtis, 2022; Spitzer & Blankenburg, 2012; Vergara et al., 2016) and/or become categorically biased (e.g., Bae, 2021; Wolff et al., 2020; Yu et al., 2020). In humans, these studies used functional imaging (fMRI), which lacks the temporal resolution to disclose rapid format changes, or EEG/MEG, which often can decode the task-relevant stimulus information only during the first 1-2 seconds of unfilled WM delays (Bae, 2021; Barbosa et al., 2021; King et al., 2016; Spitzer & Blankenburg, 2011; Wolff et al., 2017). Here, we used a different approach which leverages the finding that subtle ocular activity (e.g., microsaccades; Engbert & Kliegl, 2003; Liu et al., 2022) can reflect attentional orienting during visuospatial WM tasks (e.g., van Ede et al., 2019). While traditionally considered a confound that experimenters seek to avoid, small gaze shifts can reflect certain types of visuospatial WM information with even greater fidelity than EEG/MEG recordings (Mostert et al., 2018; Quax et al., 2019), and even throughout prolonged WM delays, which opens new avenues for tracking dynamic format changes.

Based on previous behavioral and theoretical work, we hypothesized that the level of abstraction in WM may change when the to-be-maintained information has been temporarily unattended. While unattended, WM contents cannot easily be decoded with neuroimaging approaches (but see Christophel et al., 2018), and the neural substrates of unattended storage remain disputed (Barbosa et al., 2021; Beukers et al., 2021; Stokes, 2015). Behaviorally, however, temporary inattention renders working memories less precise (Bae & Luck, 2018; Emrich et al., 2017) and more categorically biased (Bae & Luck, 2019), which may suggest increased abstraction of the WM content (see also Kerrén et al., 2022). Physiological evidence for when and how such modifications may occur during WM maintenance is still lacking.

Here, we recorded eye movements while participants memorized the orientations of rotated objects in a dual retro-cue task (Fig 1a). With such a task layout, it is commonly assumed that the initially uncued information is unattended (or deprioritized) in WM, while the cued information is in the focus of attention (Lewis-Peacock & Postle, 2012; Rose et al., 2016; Wolff et al., 2017). Representational geometry analyses borrowed from neuroimaging (Kriegeskorte & Kievit, 2013) allowed us to track with high temporal precision whether orientation encoding in gaze patterns was object-specific (indicating a concrete-visual memory), object-independent (indicating more generalized/abstract task coordinates), and/or categorically biased, throughout the different stages of the task. Participants were encouraged to keep fixation through online feedback (closed-loop) in order to restrict eye-movements to small and involuntary gaze shifts.

**Figure 1.**
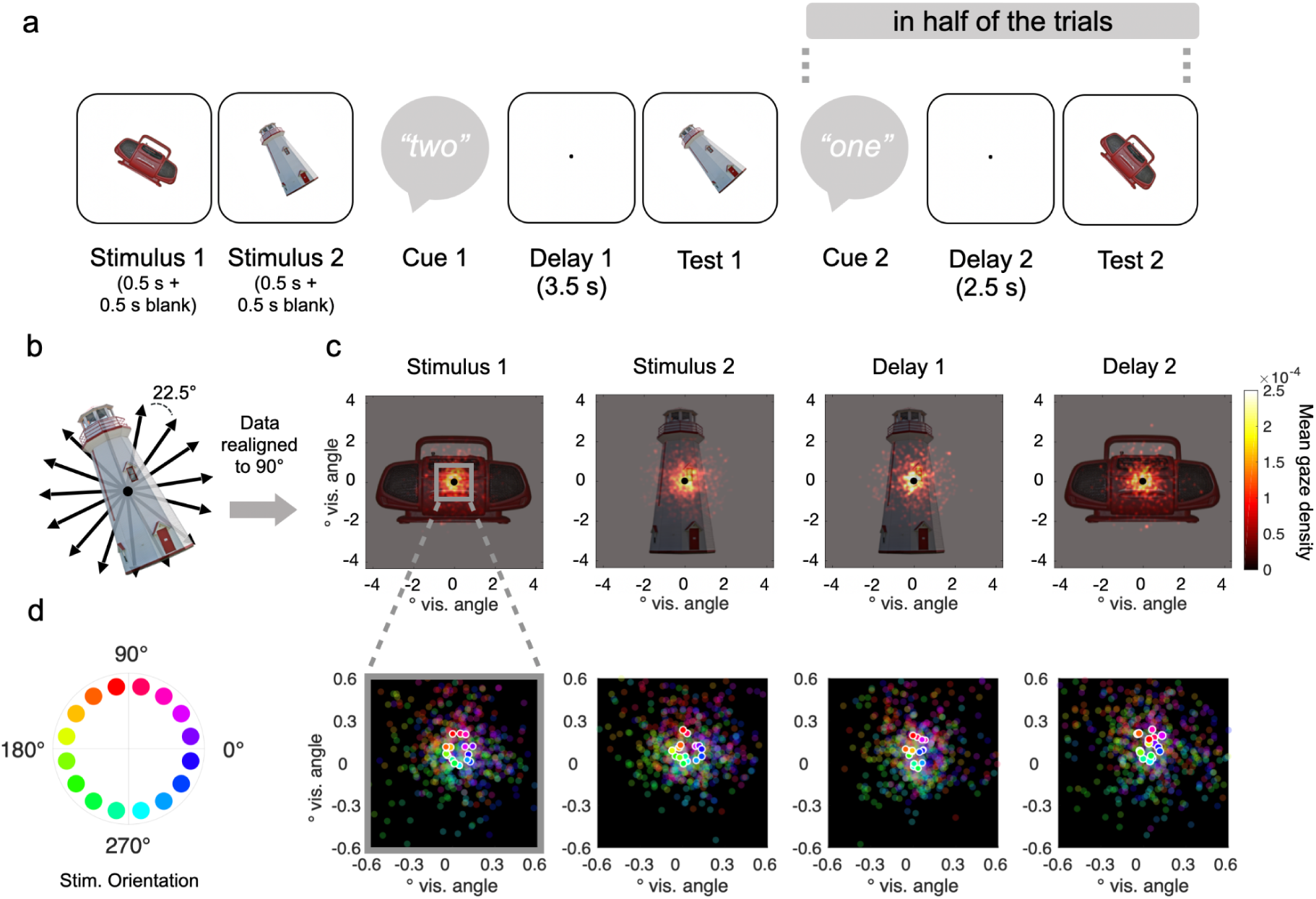
Eye-tracking during working memory for visual object orientation. **a**, Example trial. After presentation of two randomly oriented objects (Stimulus 1 and 2) an auditory cue indicated which of the two stimulus orientations was to be remembered after an unfilled retention interval (Delay 1). At test, participants were asked to re-rotate the probe stimulus to its memorized orientation (2-AFC). On half of the trials (randomized), participants were subsequently cued to also remember the orientation of previously uncued stimulus after another retention period (Delay 2 and Test 2). Participants were instructed to fixate a centered dot throughout the delay periods, and fixation breaks were penalized with closed-loop feedback from online eye-tracking (see *Methods*). **b,** Across trials, the orientations of the memory items were randomly varied around the full circle (in 16 steps, excluding cardinal orientations). **c,** Illustration of the gaze data relative to visual stimulus size. Heatmaps show gaze densities (aggregated across subjects) after aligning (re-rotating) the data from each trial to the object’s upright (90°) position. Panels show the densities aggregated over different trial periods (see *a*), with rotational alignment to the currently relevant stimulus orientation (see corresponding example objects in *a*). **d**, Mean gaze positions (without rotational alignment) during the trial periods in *c*, for the 16 different orientations of the currently relevant stimulus (see color legend in *left*). Plots show magnification (5x) of the central display area outlined in *c, left*. Saturated dots, mean; unsaturated dots, individual participants.

We found that despite this fixation monitoring, miniature gaze patterns clearly encoded the cued stimulus orientation throughout the WM delays. While the orientation encoding was object-specific at first (indicating attentional focusing on concrete visual details), its format rapidly became object-independent (generalized/abstract) when another stimulus was encoded or maintained in the focus of attention. We further found that temporary inattention increased repulsive cardinal bias in subsequent memory reports, with some evidence for such biases emerging already during the delay periods in the geometry of gaze patterns. Together, our findings indicate adaptive format changes during WM maintenance within and outside the focus of attention and highlight the utility of detailed gaze analysis for future work.

## Results

Participants (n = 41) performed a cued visual working memory task (Fig. 1a) while their gaze position was tracked. On each trial, two randomly oriented stimuli (pictures of real-world objects) were sequentially presented (each for 0.5 s followed by a 0.5 s blank screen), after which an auditory “retro”-cue (Cue 1) indicated which of the two orientations was to be remembered after a delay period (Delay 1, 3.5 s) at Test 1. On half of the trials (randomly varied), Test 1 was followed by another retro-cue (Cue 2) and another delay period (Delay 2, 2.5 s), after which participants were required to also remember the orientation of the other, previously uncued stimulus (Test 2).

### Behavioral accuracy

At each of the two memory tests, the probed stimulus was shown with a slightly altered orientation (+/-6.43°), and participants were asked to re-rotate it to its previous orientation via button press (two-alternative forced choice, 2-AFC). As expected, the percentage of correct responses was descriptively higher on Test 1 (M = 73.41%; SD = 6.42%) than on Test 2 (M = 66.62%; SD = 5.78%). Further, the second presented orientation (Stimulus 2) was remembered better (M = 70.99%, SD = 7.07%) than the first presented orientation (Stimulus 1; M = 69.04%; SD = 6.79%). A 2 x 2 repeated measures ANOVA with the factors Test (1/2) and Stimulus (1/2) confirmed that both these effects were significant [F(1,40) = 95.396, p < 0.001, η^2^ = 0.521 and F(1,40) = 13.319, p < 0.001, η^2^ = 0.043], while there was no significant interaction between the two factors [F(1,40) = 3.681, p = 0.062, η^2^ = 0.008].

The effects of presentation- and testing order may both be attributed to task periods since stimulus presentation during which the other stimulus was to be attended, either for perceptual processing (Stimulus 2) or for cued maintenance and reporting (Delay 1 and Test 1). We may combine these two factors into the “mnemonic distance” of a stimulus, which in our experiment had 4 levels (from shortest to longest: Stimulus 2 at Test 1, Stimulus 1 at Test 1, Stimulus 2 at Test 2, and Stimulus 1 at Test 2). The behavioral accuracy results were compactly described as a monotonic decrease across these distance levels [t(40) = -9.404, d = -1.469, p < 0.001; t-test of linear slope against zero], as would be expected if the processing of Stimulus 2 temporarily withdrew attention from the memory of Stimulus 1, similar (and additive) to the withdrawal of attention from the uncued item during Delay 1 and Test 1.

### Eye-tracking Results

We informed participants that their gaze would be monitored in order to ensure that they constantly fixated a centrally presented dot throughout the task. To enforce this, we provided real-time feedback (closed-loop) when fixation was lost (see *Methods*). Figure 1c shows the participants’ gaze distribution in relation to stimulus size after rotating the trial data to the respective object’s real-world (upright) orientation (for similar approaches, see Ester et al., 2016; Kang & Spitzer, 2021). Despite this rotation alignment, the gaze density was concentrated narrowly (mostly within < 1° visual angle) at center, during both the stimulus- and the delay periods (Fig. 1c). The instructions and online feedback thus proved effective in preventing participants from overtly gazing at the location of the objects’ peripheral features (such as, e.g., the spire of the lighthouse in Fig. 1a). However, inspecting the participants’ average gaze positions for each stimulus orientation (without rotational alignment) disclosed miniature circle-like patterns (Fig. 1d), indicating that miniscule gaze shifts near fixation did carry information about the objects’ orientation (for related findings with other stimulus materials, see e.g., Mostert et al., 2018; Quax et al., 2019).

#### Object orientation was robustly reflected in miniature gaze patterns

For quantitative analysis of the orientation encoding in gaze, we used an approach based on Representational Similarity Analysis (RSA; Kriegeskorte & Kievit, 2013). Specifically, we examined to what extent the gaze patterns showed the characteristic Euclidean distance structure of evenly spaced points on a circle (Fig. 2a). We implemented RSA on the single-trial level (Fig 2b) by correlating for each trial the model-predicted distances with the vector of gaze distances between the current trial and the trial average for each stimulus orientation. The procedure yields a cross-validated estimate of orientation encoding at each time point for every trial (see *Methods*).

**Figure 2.**
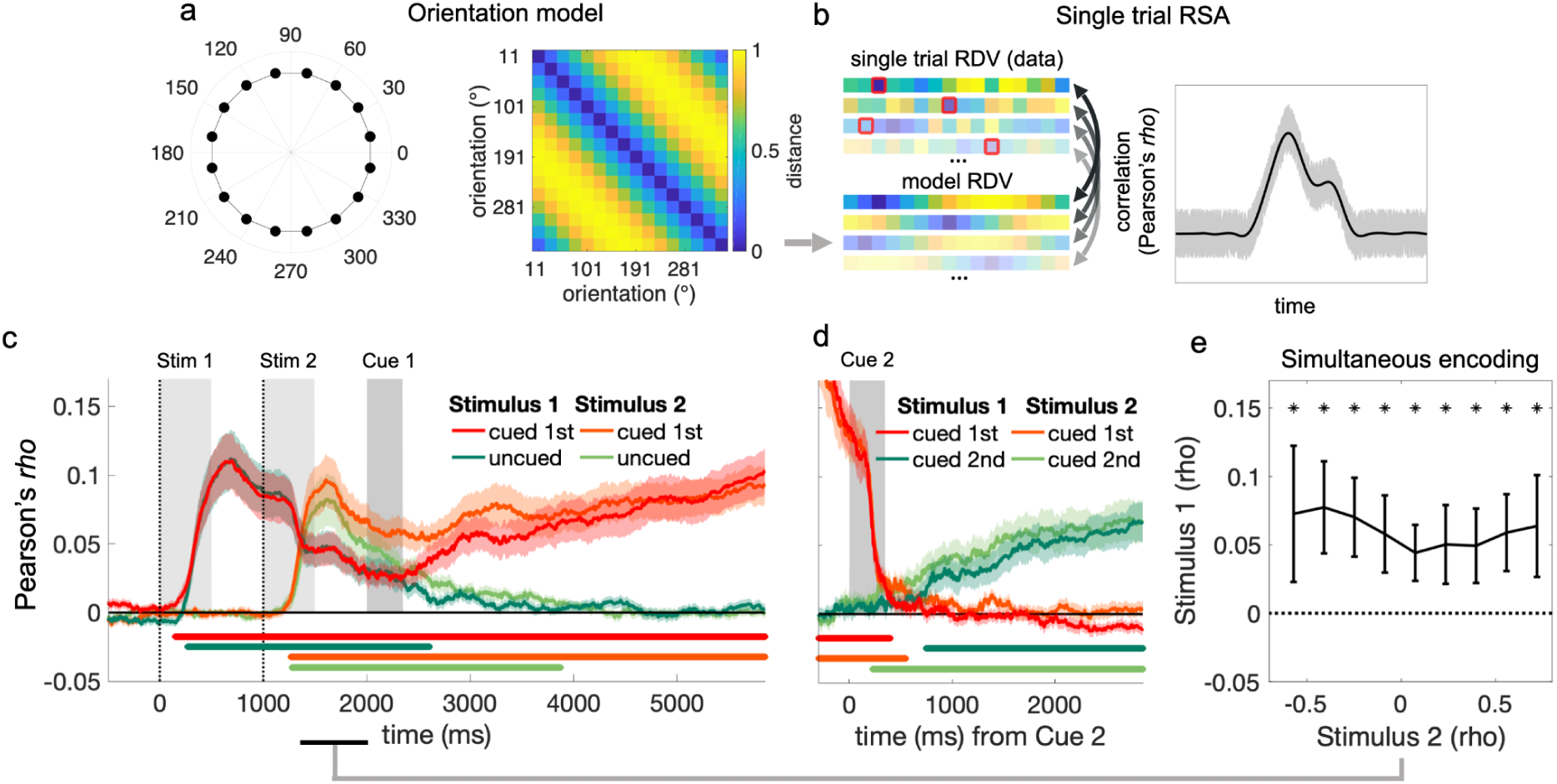
Single-trial RSA and orientation encoding time courses. **a**, *Left*: Stimulus orientations (see Fig. 1b) plotted as points on the unit circle. *Right*: Pairwise Euclidean distances between the circle points in *a*. **b**, RSA was performed at the single trial level. *Left*: Representational dissimilarity vector (RDV) of the Euclidean distances between the gaze position on the current trial (current orientation highlighted by red squares) and the mean gaze positions associated with the 16 different orientations (trial averages with the current trial data held out). Gaze RDVs were obtained at each time point and correlated trial-by-trial with the corresponding model RDV (i.e., the distances predicted by the circular orientation model in *a*), yielding a time course of orientation encoding for each trial (see *Methods*). *Right*, the single-trial approach yields a mean time-course of orientation encoding as would be obtained with conventional RSA (black lines) while additionally retaining trial-by-trial variability (gray shadings). **c**, Mean encoding of the two stimulus orientations, shown separately for when Stimulus 1 or Stimulus 2 was cued for Test 1 (see Fig. 1a). Vertical dotted lines indicate the times of stimulus onset. Gray vertical bars indicate duration of stimuli and auditory cues (“one”/”two”). Colored shadings show SEM. Colored marker lines at the bottom indicate significant orientation encoding (display threshold p_cluster_ < 0.0125 to account for testing in 4 conditions). **d**, Same as *c*, for the second delay period (Delay 2, see Fig. 1a, *right*). Note that the strong pre-cue effect is explained by residual eye-movements related to Test 1. **e**, Single-trial analysis of Stimulus 1 orientation encoding concurrent to encoding the orientation of Stimulus 2 (see time window outlined by black marker in *c* at the bottom). Trials were binned according to orientation encoding strength for Stimulus 2 (x-axis; mean values of bins averaged over participants), with orientation encoding strength for Stimulus 1 plotted on the y-axis. Error bars show SEM. Asterisks indicate significant differences from 0 (p < 0.05, Bonferroni-corrected). Significant encoding of the orientation of Stimulus 1 was evident at each bin, indicating that gaze position carried information about both stimulus orientations simultaneously.

We first inspected the mean time courses of orientation encoding (averaged over trials) during stimulus presentation. We observed robust encoding of stimulus orientation from approximately 500 ms after stimulus onset, both for Stimulus 1 and for Stimulus 2 (both p_cluster_ < 0.001, cluster-based permutation tests, see *Methods*). The encoding of either stimulus orientation peaked at approx. 650 ms (i.e., only after the stimuli’s offset; Fig. 2c), after which it slowly decayed.

#### Concurrent encoding of Stimulus 1 during encoding of Stimulus 2

Although gaze data are only two-dimensional, we found that while encoding the second presented orientation (Stimulus 2), the gaze pattern also continued to carry information about the first-presented orientation (Stimulus 1; see Fig. 2c). Such a concurrency in the average time courses may have arisen if one of the orientations was encoded on some trials and the other orientation on others. Alternatively, however, the pattern may indicate that gaze encoded both orientations simultaneously (i.e., additively, on the same trials). To shed light on this, we capitalized on our single-trial approach (see *Methods* and Fig. 2b) and binned each participant’s trials according to how strongly the orientation of Stimulus 2 was encoded between 250 and 1000 ms after Stimulus 2 onset (Fig. 2e). If encoding of the two orientations had alternated between different trials, we would expect a negative relationship with the encoding of the orientation of Stimulus 1 in the same time window. However, we found no significant relationship [t(40)= -1.368, d = -0.214, p = 0.18, linear trend analysis]. What is more, the encoding of Stimulus 1’s orientation was significantly above chance even on those trials on which the encoding of Stimulus 2’s orientation was maximally strong [t(40) = 3.656, d = 0.571, p = 0.01; t-test against 0]. Together, these results suggest that small shifts in 2D gaze space carried information about the two stimulus orientations simultaneously, on the same trials.

#### Gaze patterns encoded the cued orientations throughout the delay periods

Our main interest was in how gaze patterns reflected information storage during the unfilled delay periods (Delay 1 and 2; see Fig 1a). During Delay 1, approximately 500 ms after auditory cueing (Cue 1), the encoding of the cued orientation ramped up and continuously increased in strength until the time of Test 1, whereas the encoding of the uncued orientation slowly returned to baseline (Fig 2c). During Delay 2 (which occurred in half of the trials), a similar ramping-up pattern was observed for the second-cued orientation (which was previously uncued; Fig. 2d; both p_cluster_ < 0.001). Thus, miniature gaze deflections robustly encoded the currently cued (or “attended”) memory information during the two delay periods, in a ramp-up fashion that resembled the encoding of WM information in neural recordings (e.g., in monkey prefrontal cortex; Barak et al., 2010; Watanabe & Funahashi, 2007) .

#### Object-specific vs. object-independent orientation encoding

We next examined more closely in which format(s) the gaze patterns reflected the WM information. A priori, memory reports in our task could be based on a concrete visual memory of the presented stimulus, but they could also be based on a mental abstraction of orientation, e.g., in terms of directional spatial coordinates. To the extent that the small eye movements during WM maintenance reflected mental focusing on concrete visual features (e.g., the location of a specific point on the object’s contour), we expect the orientation encoding in gaze to be object-*specific,* that is, not fully transferable between different objects. In contrast, an abstraction of orientation (e.g., in terms of a direction in which any object may point with its real-world top) should be reflected in gaze patterns that are object-*independent* and transferable.

We examined object-specificity by comparing the orientation encoding in gaze distances *within* objects (Fig. 3a, *left*) with that in gaze distances *between* objects (Fig. 3a, *right*). Upon stimulus presentation, the orientation encoding in gaze patterns was object-specific, in that within-objects encoding clearly exceeded between-objects encoding (Fig. 3c, all p_cluster_ < 0.012). For Stimulus 1, the object-specificity diminished abruptly after approx. 1300 ms (when the gaze patterns began to also encode Stimulus 2), and changed to a more object-independent format for the remainder of the trial epoch (Fig. 3c, *upper*). For Stimulus 2, in contrast, when cued for Test 1, the object-specificity decayed less and was sustained throughout most of Delay 1 (Fig. 3c, *lower*). Later, in Delay 2, no object-specificity was evident anymore for either stimulus (no p_cluster_ < 0.60). Figure 3e-f summarize the temporal evolution of object-independence in terms of Bayes Factors, showing the swift change of Stimulus 1 encoding from object-specific (BF_01_ < ⅓) towards object-independent (BF_01_ > 3) at the time of Stimulus 2 encoding, whereas the encoding of Stimulus 2 retained object-specificity during Delay 1 (Fig. 3e). In Delay 2, after unattended storage throughout Delay 1, the orientation encoding in gaze had become object-independent for both stimuli (Fig. 3f).

**Figure 3.**
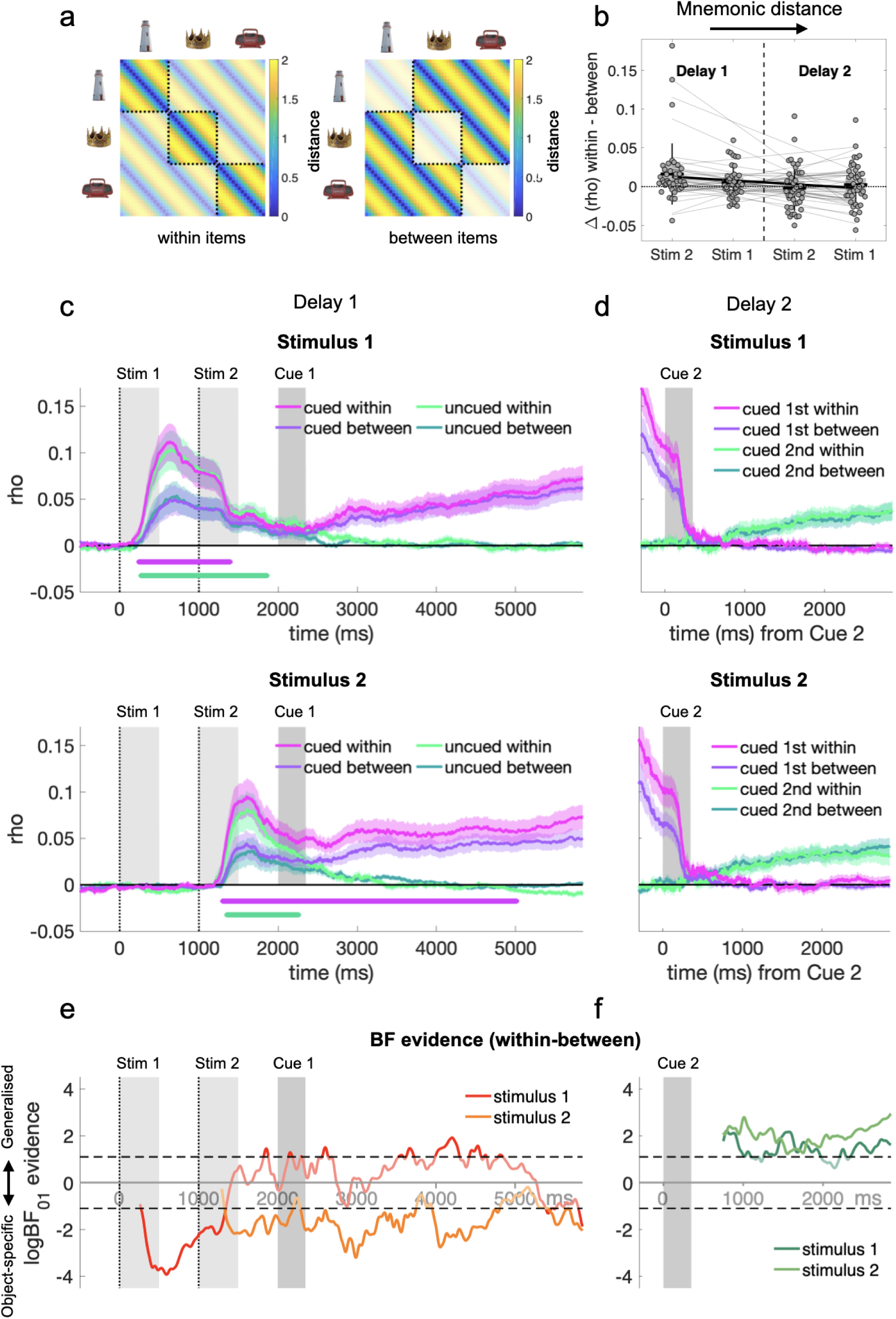
Object-specific vs. object-independent orientation encoding. **a**, Orientation model analogous to Fig. 2a, but extended to separately examine orientation encoding within and between objects. *Left,* within-objects orientation encoding; *Right,* between-objects orientation encoding. Unsaturated colors delineate distances that are excluded from the respective sub-model. The degree of object-specificity is inferred from the extent to which within-objects encoding is stronger than between-objects encoding. **b**, Difference in orientation encoding within objects compared to between objects, for the cued stimulus during each delay period. Data are averaged from cue onset to the end of the delay. The four conditions are sorted by the time distance from stimulus presentation (“mnemonic distance”, from lowest to highest). Gray dots show individual participants results and trend lines show linear fits. Box plots show group means with SEM (boxes) and SD (vertical lines). **c**, Orientation encoding time courses within and between objects during Delay 1, shown separately for when the stimulus was cued (pink- and purple) or uncued (light and dark green). *Top*: Stimulus 1; *bottom*: Stimulus 2. Colored shadings show SEM. Colored marker lines on the bottom indicate significant object specificity in terms of stronger orientation encoding within than between objects (display threshold p_cluster_ < 0.0125). Otherwise same conventions as Fig 2c. **d**, Same as *c*, but for the second delay period (Delay 2). **e-f**, Bayes Factor (BF_01_, one-tailed) analysis of the difference between within- and between-objects encoding. BF time courses are shown for the cued orientation in the respective task periods. Negative values on the log scale (y-axis) indicate stronger evidence for object-specific encoding (within > between) than for object-independent encoding (within ≤ between), positive values indicate the opposite. Results are shown for periods of significant overall orientation encoding (cf. Fg. 2) where the comparison of within- and between objects encoding is meaningful. The data were smoothed with a 50 ms Gaussian kernel before this analysis. Saturated colors indicate stronger than anecdotal evidence (logBF < -1.1 or > 1.1, which corresponds to BFs <1/3 or > 3).

Focusing on the delay periods, we examined whether the object-specificity of cued orientation encoding differed between the two delay periods (Delay 1 or 2) and/or between the first and second presented stimulus (Stimulus 1 or 2). A 2 x 2 repeated-measures ANOVA on the difference in encoding strength (within-minus between-objects, averaged across the respective delay periods) showed a main effect of delay period [Delay 1/2; F(1,40) = 6.204, p = 0.017, η^2^ = 0.064], indicating greater object-specificity during Delay 1, but no effect of presentation order [Stimulus 1/2; F(1,40) = 0.985, p = 0.327]. There also was a moderate interaction between the two factors [F(1,40) = 4.466, p = 0.041, η^2^ = 0.027], reflecting that the difference between the two delays was stronger for Stimulus 2. Again, we also inspected these results in terms of the mnemonic distance from stimulus presentation, that is, the time the orientation in question had been unattended while focusing on the other orientation (Fig. 3b). Indeed, this analysis confirmed a decrease of object specificity with increasing mnemonic distance [t(40) = -2.473, d = -0.386, p = 0.018; t-test of linear slope against zero].

Together, these results showed that unlike during perceptual processing, gaze patterns during the delay periods reflected WM information in more generalized (or abstract) coordinates, and that the level of this abstraction increased after periods of temporary (or partial) inattention.

### Cardinal repulsion bias in gaze patterns and behavior

In studies of WM for stimulus orientation (e.g., of Gabor gratings) it is commonly observed that behavioral reports are biased away from the cardinal (vertical and horizontal) axes (Bae, 2021; Taylor & Bays, 2018). We asked (i) whether such a repulsive cardinal bias also occurred with our rotated object stimuli, (ii) whether the strength of bias was modulated by periods of inattention (Bae & Luck, 2019), and (iii) to what extent such bias was already expressed in the geometry of the miniature gaze patterns observed during the delay periods.

To model bias in behavior, we used a geometrical approach that quantifies bias as a mixture of a perfect (unbiased) circle (Fig. 4a, *middle*) with perfect (fully biased) square geometries (Fig. 4a, *leftmost* and *rightmost*, see *Methods, Behavioral modeling* for details). Intuitively, the mixture parameter 𝐵 quantifies the extent to which the reported orientations were repulsed away from the cardinal axes, with 𝐵 > 0 indicating repulsion (i.e., cardinal bias), 𝐵 = 0 no bias, and 𝐵 < 0 attraction.

**Figure 4.**
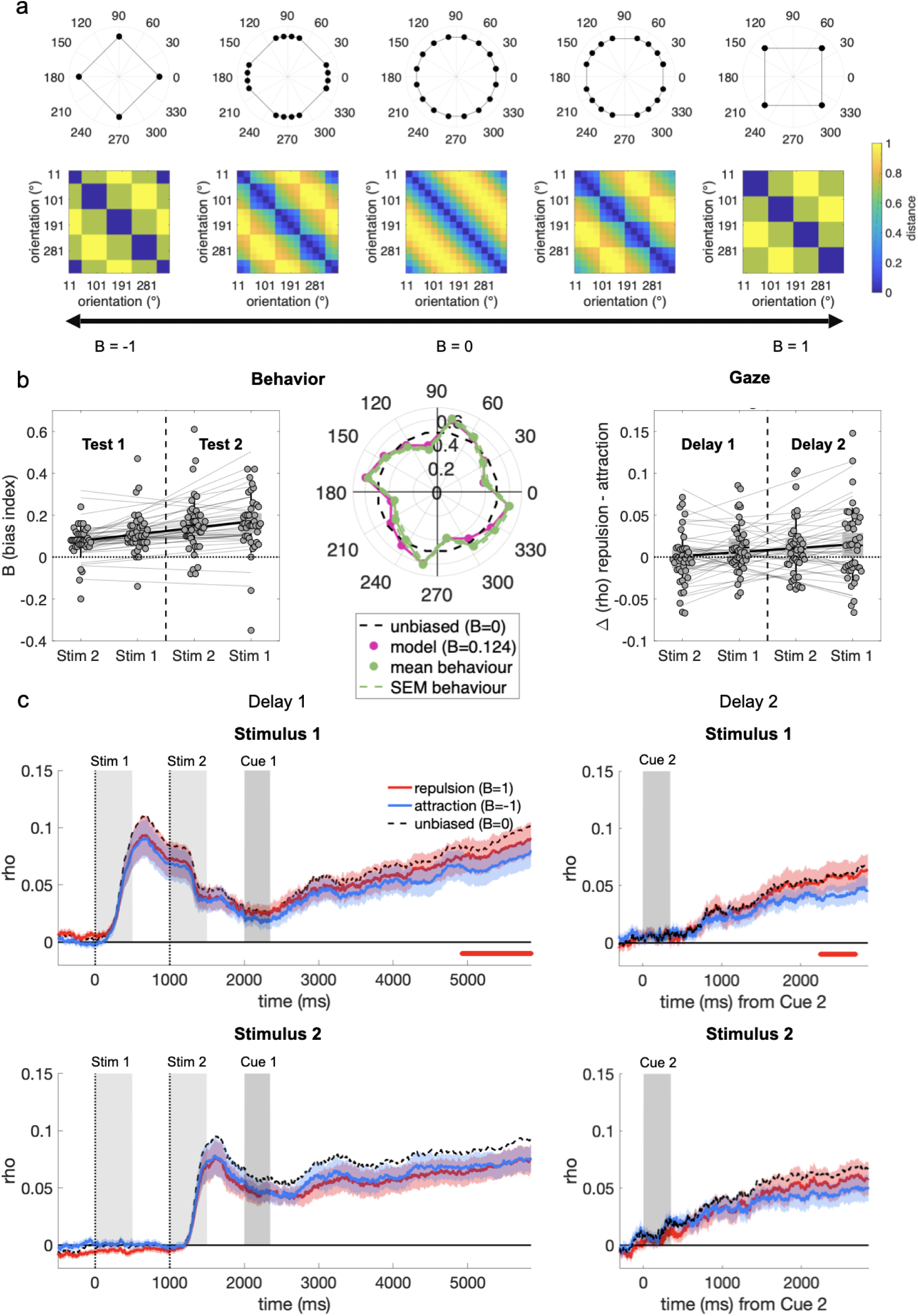
Cardinal repulsion bias in gaze patterns and behavior. **a**, Model-predicted geometries (top) and RDMs (bottom) associated with different levels of cardinal bias. The mixture parameter 𝐵 (black arrow axis on bottom) denotes the level of repulsion (𝐵 > 1; “cardinal bias”) or attraction (𝐵 < 1) from/to cardinal axes, relative to the unbiased circle model (𝐵 = 0). **b**, *Left and Middle:* Results from the model fitted to the behavioral memory reports *Left*: Estimates of cardinal repulsion bias (𝐵) for each stimulus (1/2) and test (1/2), sorted by the distance between stimulus presentation and test (same conventions as Figure 3b). *Middle*: Polar plot shows the mean proportions of clockwise reports (green) and the predictions of the fitted model (magenta; 𝐵 = 0.12) for each stimulus orientation. Results are averaged over both stimuli and tests. Dashed black line shows proportions (50%) expected under an unbiased circular model (𝐵 = 0) for visual reference. *Right*: Quantification of bias in the gaze patterns associated with the cued stimulus orientation. Shown are the differences in correlation of the gaze patterns with the repulsion model (𝐵 = 1) compared to the attraction model (𝐵 = -1), where a positive difference indicates repulsive cardinal bias. Results are shown for the last second of the respective delay period (see *c*). **c,** Time course of correlations with the repulsion (red) and attraction (blue) models during the delay periods (same layout as Fig. 3c-d). Colored shadings show SEM. Dashed black line shows correlation with the unbiased circular model (𝐵 = 0) for visual reference. Red marker lines on the bottom indicate stronger correlation with the repulsion than the attraction model (display threshold p_cluster_ < 0.05).

Fitting the model to participants’ behavioral responses (Fig. 4b, *left* and *middle*), we observed values of 𝐵 > 0 (grand mean: 0.124, SD = 0.092) in both memory tests (Test 1 and 2) and for both orientations [Stimulus 1 and 2; all 𝐵 > 0.071; all t(40) > 4, all p < 0.001; t-tests against 0]. Thus, participants overall showed a repulsive cardinal bias, which replicates and extends previous work with simpler stimuli (such as gratings; Bae, 2021; Taylor & Bays, 2018). A 2 x 2 repeated measures ANOVA showed a main effect of Test [1/2; F(1,40) = 19.743, p < 0.001, η^2^ = 0.144], indicating a stronger bias on Test 2, and a main effect of presentation order [Stimulus 1/2; F(1,40) = 4.669, p = 0.037, η^2^ = 0.024] with no interaction between the two factors [F(1,40) = 1.083, p = 0.304]. The overall pattern could again be described compactly as an increase in cardinal bias with increasing mnemonic distance from stimulus presentation [t(40) = 5.315, d = 0.830, p < 0.001; t-test of linear slope against zero, Fig. 4b, *left*]. Thus, we found robust cardinal repulsion in participants’ overt memory reports, and this bias increased with periods of unattended storage.

Finally, we addressed to what extent the cardinal bias was also reflected in the gaze patterns recorded throughout the two delay periods. To do so, our geometric model yields distinctive distance structures for extreme cardinal repulsion (𝐵 = 1; Fig. 4a, *rightmost)* and attraction (𝐵 = -1; Fig. 4a, *leftmost*), respectively. If the gaze patterns were unbiased, we would expect both these ‘square’ models to correlate less well with the data than the unbiased (‘circle’) model with 𝐵 = 0 (Fig. 4a *middle*). However, to the extent that the gaze patterns were repulsively biased, we would expect the repulsion model to outperform the attraction model, nearing (or in the case of extreme bias, even exceeding) the circle model (dashed black in Fig. 4c). Contrasting repulsion and attraction models thus allowed us to quantify the extent of repulsive or attractive bias in the gaze patterns during the delay periods.

Descriptively, the three different models (repulsion, unbiased, attraction) showed only small differences in correlation with the data (Fig. 4c), indicating that the statistical power to detect bias in the gaze data was relatively low (see *Methods*, *Model geometries*). Nevertheless, contrasting the repulsion model with the attraction model showed mildly significant clusters (p_cluster_ = 0.02 and p_cluster_ = 0.035), indicating a repulsive bias near the end of the delay periods for Stimulus 1 (Fig. 4c, upper). A similar tendency for Stimulus 2 failed to reach significance in Delay 2 (Fig. 4c, lower right; p_cluster_ = 0.085, below display threshold) and was absent in Delay 1 (Fig. 4c, lower left; no cluster-forming time points). A 2 x 2 repeated measures ANOVA on the difference between repulsion and attraction models (averaged across the last second of the delay periods) showed a main effect of presentation order [Stimulus 1/2; F(1,40) = 4.561, p = 0.039 η^2^ = 0.026; main effect of Delay 1/2: F(1,40) = 1.650, p = 0.206; interaction: F(1,40) < 1], indicating a stronger repulsive bias for the first presented orientation (Stimulus 1). Complementary analysis in terms of mnemonic distance (Fig. 4b, *right*) showed a positive trend similar to that in behavior, albeit only at the significance level of a one-tailed test [t(40) = 1.772, d = 0.278, p = 0.042; t-test of linear slope against zero; one-tailed, hypothesis derived from behavioral result]. Together, while the differentiation of models (repulsive, unbiased, attractive) in the gaze data was not as clear-cut as in behavior (cf. Fig. 4b, *right* and *left*), we found indications that the gaze patterns may have carried a repulsive cardinal bias, most evidently during the later portions of the WM delays and after temporary and/or partial inattention to the WM information.

### Orientation-dependent microsaccades

While our RSA-based approach was designed to characterize the time-varying geometries of aggregate gaze position patterns (Figs 2-4), we performed additional analysis on request by reviewers to explore whether the findings may indeed be related to microsaccades (see *Introduction*), i.e., “jerk-like” (Rolfs, 2009) small eye movements. Figure 5 illustrates the directions of microsaccades detected after stimulus presentation (Stimulus 1, Stimulus 2) and during the two delay periods (Delay 1, Delay 2), respectively, for each of the 16 orientations of the currently relevant object. The saccade directions in the post-stimulus periods correlated positively with stimulus orientation (circular correlation: Stimulus 1, R = 0.089; Stimulus 2, R = 0.077; both p < 0.001). Weakly positive correlations were also evident during the delay periods (Delay 1, R = 0.01; Delay 2, R = 0.03; both p < 0.001). For further inspection, we again rotated the trial data (analogous to Fig. 1b-c) to illustrate the saccade directions relative to the objects’ real-world (upright) orientation. As expected if microsaccades reflected stimulus orientation, the aligned distributions were not uniform (all z’s > 34.36, all p < 0.001, Rayleigh tests for uniformity) but appeared egg-shaped, with a main peak near the object’s real-world top (at 90°) and another, smaller peak near the opposite angle (270°, which may reflect ‘return’ microsaccades to fixation). Together, these complementary results support the idea that the effects observed in our main analyses may have been related to microsaccadic activity during attempted fixation (Engbert & Kliegl, 2003; Hafed & Clark, 2002).

**Figure 5.**
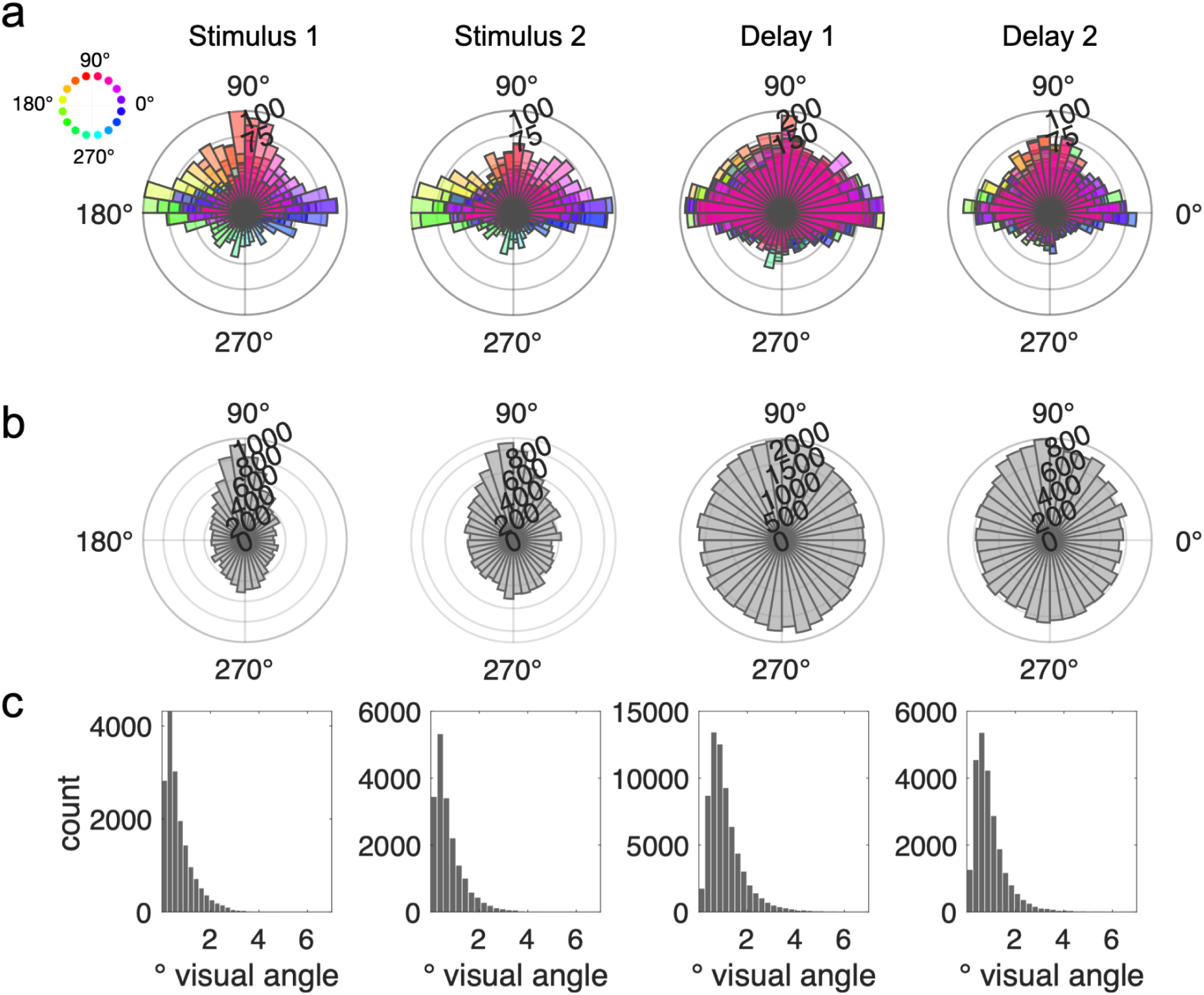
Complementary analysis of microsaccadic activity. We performed microsaccade detection (see *Methods*) in four time windows during which orientation encoding was evident in the gaze position data (see Fig. 2c-d): from 300 to 1000 ms after the onset of each stimulus (Stimulus 1, Stimulus 2), from after Cue 1 until the end of the first delay (Delay 1), and from 300 ms after Cue 2 until the end of the second delay (Delay 2). **a**, Distribution of microsaccade directions (onset to endpoint, see *Methods*) as a function of stimulus orientation (see color legend on the left), collapsed across all trials from all participants. The distributions during the delay periods (Delay 1 and 2) are color-coded according to the currently cued orientation, respectively. **b**, Distribution of microsaccade directions after alignment (analogous to Fig. 1b-c, upper) relative to the objects’ upright (90°) position. **c**, Histograms of the sizes (amplitudes) of the saccades detected for analysis in *a* and *b*. The majority of saccades in each of the time windows was considerably smaller than 2°, in line with an interpretation in terms of microsaccadic activity (Rolfs, 2009).

## Discussion

The processing of WM information during delay periods has been studied extensively using neural recordings (for reviews, see Christophel et al., 2017; D’Esposito & Postle, 2015; Goldman-Rakic, 1995; Miller et al., 2018; Stokes, 2015; Wang, 2021). Here, using novel stimulus materials and tailored geometry analyses, we showed that miniature gaze deflections can disclose an array of WM-associated phenomena that were previously only observed in neural signals, including (i) a sustained encoding of the task-relevant stimulus feature, which (ii) shows a different format than during perception, can (iii) persist while also encoding new perceptual information, (iv) ramps up throughout delay periods when relevant for an upcoming test, and (v) returns to baseline when uncued (or “unattended”). Beyond this, the gaze geometries indicated that temporary inattention rendered the WM information more generalized (object-independent), and potentially more categorically biased. These format changes during maintenance were similarly observed when attention to the memorandum was withdrawn by explicit retro-cueing or by presenting additional WM information.

Behaviorally, our results replicate and extend previous findings that temporary inattention renders working memories less precise and more biased (Bae & Luck, 2019; Emrich et al., 2017). The eye-tracking results shed light on the temporal unfolding of potentially underlying format changes during WM storage. The gaze patterns during perceptual processing were clearly object-specific, indicating a focus on concrete visual details. When the last seen stimulus (Stimulus 2) was immediately cued (with an auditory retro-cue that offered no visual distraction), some of this object-specificity was sustained throughout the ensuing WM delay. In contrast, for the first-presented stimulus (Stimulus 1), the object-specificity dropped abruptly as soon as Stimulus 2 processing commenced. Of note, Stimulus 1 encoding did continue throughout Stimulus 2 processing. However, its format changed to object-independent (or “abstract”) during the object-specific (or “concrete”) encoding of Stimulus 2 — as if the memory of Stimulus 1 was reformatted to ‘evade’ the format of the currently perceived stimulus. Subsequent cueing did not revert this effect, nor did we find any re-emergence of object-specificity after unattended storage, for either stimulus, in the second delay. Together, these results support the idea that temporary (or partial) inattention may render the task-relevant WM information (here, orientation) increasingly less “concrete” (visual-sensory) and more generalized or “abstract”.

Further support for this idea comes from our analysis of the cardinal repulsion bias. In parallel with the object-independence of gaze patterns, the repulsive cardinal reporting bias increased with the time a given stimulus had been temporarily (or partially) unattended. Repulsive orientation bias in WM tasks has been explained, e.g., by efficient coding principles, in terms of relatively finer tuning to cardinal orientations, reflecting their relative prevalence in natural environments (Taylor & Bays, 2018). An alternative framing of the cardinal repulsion biases in our experiment with real-life objects could be in terms of more explicit semantic categorization (e.g., “left”/“right” and “up”/“upside down”; Hardman et al., 2017; Ricker et al., 2022). The results may thus also reflect increased reliance on semantics (Kerrén et al., 2022) and/or (pre-)verbal labels when restoring information from unattended storage (see also Beukers et al., 2021), which would be in line with a higher level of abstraction. While our geometrical analysis approach is agnostic to the mechanistic cause of cardinal biases, we found some indications that they were also evident in biased gaze geometries during the stimulus-free retention periods (for related findings in neuroimaging, see Bae, 2021; Ester et al., 2020; Wolff et al., 2020). The latter result was statistically weak and should be revisited in future work, possibly under conditions that induce even stronger biases in behavior (e.g., higher WM loads; Taylor & Bays, 2018).

A remarkable aspect of our results is the small amplitude of the eye-movements that disclosed such rich information. The mass of the raw position samples in our analysis was within < 1° visual angle around fixation (Fig. 1c). A discernible “circular” structure in averaged data points (Fig. 1d) measured only approx. 0.2-0.3° in diameter, which is near the eye-tracker’s accuracy limit, and was only a fraction of the memory items’ physical size. Together with our online fixation control (see Methods), these descriptives render it unlikely that our results were attributable to reflexive saccades to the location of peripheral stimulus features. Additional analysis (Fig. 5) indicated that the findings more likely reflect microsaccadic activity during attempted fixation. Systematic microsaccade patterns have previously been linked to covert spatial attention (Engbert & Kliegl, 2003; Hafed & Clark, 2002), suggesting that in the present context, they might have reflected mental orienting towards a spatial coordinate or direction (Liu et al., 2022; van Ede et al., 2019). Together, our results indicate that participants generally oriented attention towards the objects’ real-life “top”, but with varying degrees of bias towards specific object features (resulting in object-specific orientation patterns), and/or away from cardinal axes (resulting in cardinal repulsion).

Under a view of the miniature gaze patterns reflecting covert spatial attention, our analysis tracked with high temporal resolution the time course of attention allocation to WM information in a dual retro-cue task. Before cueing, encoding a new stimulus (Stimulus 2) did not immediately eradicate or replace the attentional orienting to the previous stimulus (but did change its qualitative format, see above). At face value, the temporary simultaneity of both WM contents (Fig. 2e), in a putative index of attention, might appear to be at odds with the idea of an exclusive single-item focus of attention in WM (Oberauer, 2002; Olivers et al., 2011; but see Beck et al., 2012; Zhang et al., 2018). However, another possible interpretation is that the (re)allocation of attention to different stimuli (or tasks) in WM may take time to complete. For instance, the encoding of the uncued stimulus fully returned to baseline only approx. 0.5-1.5 s after the cue, which is broadly consistent with previous behavioral and EEG work on the time course of WM-cueing effects (LaRocque et al., 2013; Souza & Oberauer, 2016; Spitzer et al., 2014; Spitzer & Blankenburg, 2011). Compared to this, the reformatting into a more generic, object-independent format was rapid, both for Stimulus 1 when encoding Stimulus 2 (see above), and for Stimulus 2 itself when it was uncued. Consistent with these results, a recent study found that low-level perceptual bias induced by concurrent WM information (cf. Teng & Kravitz, 2019) dissipated quickly with new visual input (Kang & Spitzer, 2021). These findings are in line with adaptive format changes in WM, potentially providing fast protection from interference beyond the overall reallocation of attention between different stimuli and/or tasks.

Previous WM studies using retro-cues yielded mixed results regarding potential costs for the first of two successively presented stimuli. Using visual retro-cues, one study found no differences between visual gratings presented first or second, neither in behavior nor in fMRI-decoding during the WM delay (Harrison & Tong, 2007), which has been taken as evidence that intervening stimuli may cause little to no interference for visual WM representations (Rademaker et al., 2019). Another study, using visual retro-cues with tactile WM stimuli, did find lower performance for the first stimulus (Spitzer and Blankenburg, 2011), a finding we replicated here with auditory cueing of visual WM information. One possibility is that different-modality cues (e.g., auditory when the WM stimuli were visual) interfere less with the short-term memory of the last-presented stimulus than same-modality cues would (e.g., visual cues with visual WM stimuli; for related findings, see Bae and Luck; 2019). Different-modality cues may thus leave the memory trace of the last-presented stimulus more intact compared to the first stimulus (which is always followed by the same-modality input of the second stimulus). This aside, the format changes induced by the intervening stimulus were qualitatively similar to those after unattended storage, in line with a common explanation in terms of temporarily withdrawn attention.

Our findings of increasingly more object-independent gaze geometries do not rule out that the brain may maintain detailed visual memories in ways that would not register in eye-tracking. More generally, we can only speculate whether the minuscule eye-movements observed in our experiment played a functional role or whether they were merely epiphenomena of other processes. We consider it possible that our paradigm promoted aspects of WM-related processing to become visible at the surface of ocular activity, but that the ocular activity itself may have had little or no direct role in the WM processing proper (for related discussion, see Liu et al., 2022; Loaiza & Souza, 2022; Rolfs, 2009; but see Ferreira et al., 2008; Johansson & Johansson, 2014, for a role of eye-movements in episodic memory retrieval). This speculation also takes note of several recent failures to decode visuospatial WM information from eye-tracking, most notably in control analyses supplemental to neural decoding, where systematic eye-movements were ruled out as a potential confound (e.g., Brissenden et al., 2018; Günseli et al., 2022; Kwak & Curtis, 2022; Muhle-Karbe et al., 2021; but see Mostert et al., 2018; Quax et al., 2019; Thielen et al., 2019). At the same time, our findings sound a cautionary note that stimulus-dependent eye movements in visual WM tasks can be very small, hard to prevent, persistent, and above all, informative.

In summary, despite discouraging participants from eye motion through closed-loop fixation control, we found the orientation of visual objects robustly reflected in miniature gaze patterns during cued WM maintenance. The geometry of the gaze patterns underwent systematic changes, suggesting that temporary inattention increased the level of abstraction (and categorical bias) of the information in WM. Stimulus-dependent eye movements may not only pose a potential confound, but also a valuable source of information in studying visuospatial WM.

## Methods

### Participants

Fifty-five participants (31 female, 24 male, mean age 26.95 ± 3.98 years) took part in the experiment. Forty-four of the participants were recruited from a pool of external participants and 11 were recruited internally within the Max Planck Institute for Human Development. All participants were blind to our research questions and all of them received a compensation of €10 per hour plus a bonus based on task performance (€5 bonus if four out of five randomly selected memory reports were correct). Written informed consent was obtained from all participants, and all experiments were approved by the ethics committee of the Max Planck Institute for Human Development. Two participants (both wearing glasses) were excluded due to difficulties in acquiring a stable eye-tracking signal, and one participant was excluded because she reported feeling unwell during the experimental session. Of the remaining participants, we excluded n = 9 for failing to perform above chance level in each of the two memory tests (p < 0.05, Binomial test against 50% correct responses). Finally, after preprocessing the eye-tracking data, we excluded n = 2 participants for whom more than 15% of the data had to be rejected due to blinks and other recording artifacts. After this, n = 41 participants remained for analysis.

### Stimuli, Task, and Procedure

Nine color photographs of everyday objects from the BOSS database (candelabra, table, outdoor chair, crown, radio, lighthouse, lamppost, nightstand, gazebo) were used as stimuli. All objects were cropped (i.e., background removed), and one object (gazebo) was slightly modified using GNU image manipulation software (http://www.gimp.org) to increase its mirror-symmetry. We grouped the pictures into 3 different sets of three, always combining objects with different aspect ratios (width/height; see example set in Fig. 3a). Each participant was assigned one of these sets, with each set being used similarly often across the participant sample (two sets were used 18 times and one set 19 times). As auditory cue stimuli, we prepared recordings of the words “one”, “two”, and “thanks” spoken by a female lab member. The recordings were time-compressed to a common length of 350ms using a pitch-preserving algorithm provided in Audacity® (GNU software; https://www.audacityteam.org/).

Each trial started with a fixation dot (8×8 px, corresponding to 0.17 x 0.17° visual angle) displayed at the center of the screen for 500 to 1000 ms (randomly varied), followed by sequential presentation of two objects, each in a random orientation (see below). Each stimulus was displayed for 500 ms (display size approx. 6.5° visual angle, see Fig. 1c) followed by a 500 ms blank screen. After this, an auditory retro-cue (“one” or “two”, 350 ms) indicated which of the two stimulus orientations was to be reported after a delay (Delay 1, 3500 ms) in the upcoming memory test (Test 1). Test 1 started with the cued object reappearing on display, but with its previous orientation changed by +/-6.43°. Participants were asked to indicate via key press (2-AFC) whether the object would need to be rotated clockwise or anticlockwise (right and left arrow key) to match its memorized orientation. Upon key press, the object rotated accordingly (by 6.43°), followed by a written feedback message (“correct” or “incorrect”) displayed in the upper part of the screen (500 ms). After another 500ms, in half of trials, an auditory message (“thanks”, 350 ms) signaled the end of the trial. In the other half of trials (randomly varied), a second auditory retro-cue (Cue 2) was presented (e.g., “two”, if the first retro-cue was “one”), indicating that the thus far untested stimulus orientation would still need to be reported. In these trials, another delay period ensued (Delay 2; 2500 ms), and participants’ memory for the second-cued stimulus was tested (Test 2), using the same procedure as before for the first-cued stimulus in Test 1. Each participant performed 16 blocks of 32 trials, for a total of 512 trials (265 of which included a Test 2).

Stimulus presentation was pseudo-random across trials, with the following restrictions: (i) each pairing of objects from the participant’s object set occurred equally often, (ii) each object was equally often presented first (as Stimulus 1) and second (as Stimulus 2), and (iii) Stimulus 1 and Stimulus 2 were equally often cued for Test 1. The orientations of the two objects on each trial were drawn randomly and independently from 16 equidistant values (11.25° to 348.75° in steps of 22.5°), which excluded the cardinal axes (0°, 90°, 180°, and 270°).

The experiment was run using Psychophysics Toolbox Version 3 (Brainard & Vision, 1997) and the Eyelink Toolbox (Cornelissen et al., 2002) in MATLAB 2017a (MathWorks). The visual stimuli were presented on a 60×34 cm screen with a 2560×1440 px resolution and a frame rate of 60 Hz. The auditory cue words were presented via desktop loudspeakers (Harman Kardon KH206). To minimize head motion, participants performed the experiment with their head positioned on a chin rest with a viewing distance of approx. 62cm from the screen. Gaze position was monitored and recorded throughout the experiment at a sampling rate of 500 Hz using a desktop-mounted Eyelink 1000 eye-tracker (SR research), with file and link/analog filters set to ‘EXTRA’ and ‘STD’, respectively.

Participants were instructed to constantly keep their gaze on the fixation dot, which was displayed throughout the entire trial except for the test- and feedback periods. Whenever a participant’s gaze deviated more than 71 pixels (1.53° visual angle) from the center of the fixation dot either before object presentation or for longer than 500 ms during any of the two delay periods, a warning message (“Fixate”) was displayed at the center of the screen. This occurred during less than 15% (Mean: 13.03%) of the trial epoch on average.

### Behavioral modeling

To model participants’ behavioral memory reports (2-AFC), we used a geometrical approach similar to that used in our eye-tracking analyses (see below). We first defined three prototypical geometries, (i) an unbiased “circle” model (*𝑀_circle_*) corresponding to the memory items’ 16 original orientations (see Fig. 4a, *middle*), (ii) a cardinal model (𝑀_*repulsion*_) which shifts the 16 orientations to the nearest diagonal orientation (i.e., 45°, 135° 225°, or 315°; see Fig. 4a, *rightmost*) and (iii) a cardinal model (𝑀_*attraction*_) which shifts them to the nearest cardinal orientation (i.e., 0°, 90°, 180°, or 270°; see Fig. 4a, *leftmost*). The continuum from attraction to repulsion was formalized with a mixture parameter 𝐵 (ranging from -1 to 1) which blends the circle model with the repulsion model for 𝐵 ≥ 0 :

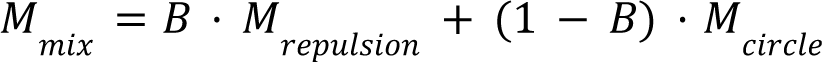

and with the attraction model for 𝐵 < 0 :

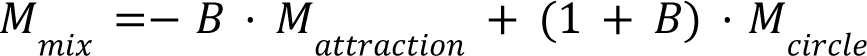

where · denotes scalar multiplication. Figure 4a illustrates the resulting model continuum from 𝐵 = -1 (maximal attraction) over 𝐵 = 0 (unbiased) to 𝐵 = 1 (maximal repulsion). To simulate memory reports (clockwise or counter-clockwise) for each trial, we computed the angular difference 𝑑 between the orientation modeled in 𝑀_*mix*_ and the probe orientation displayed at test, and transformed it into a probability of making a “clockwise” response using a logistic choice function:

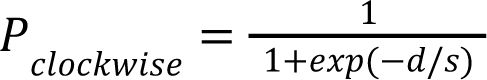

where 𝑠 is a noise parameter that relates inversely to memory strength or -precision (see also Schurgin et al., 2020). For completeness, our model also allowed for greater memory precision near the cardinal axes (a so-called “oblique effect”, e.g., Pratte et al., 2017; Taylor & Bays, 2018). This was implemented by an additional parameter 𝑐, which up- or downregulated noise 𝑠 for those 8 orientations in the stimulus set that were near to the cardinal axes (see Fig. 4a, *middle*), relative to the remaining 8 orientations that were nearer to the diagonal axes:

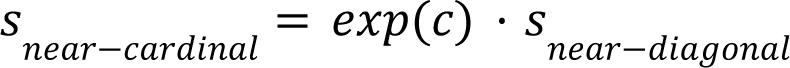

where values of 𝑐 < 0 would indicate relatively greater precision (lower noise) near the cardinal axes (i.e., an oblique effect). The model was fitted to the memory reports of each participant individually, using exhaustive gridsearch (𝐵, -1…1; 𝑠, 0…1; 𝑐, -0.5…0.5; with a step size of 0.01 for each parameter) and least squares to identify the best-fitting parameter values.

While our analysis focused on bias (𝐵), we note for completeness that we also observed values of 𝑐 significantly smaller than 0 [mean across conditions: 𝑐 = -0.292, t(40) = -10.159, p < .001, t-test against 0], i.e., an oblique effect, which replicates and extends previous work (e.g., Taylor & Bays, 2018). The strength of this oblique effect tended to decrease with mnemonic distance [t(40) = 2.286, d = 0.357, p = 0.028; t-test of linear slope against zero] (see Taylor & Bays, 2018 for related findings).

### Eye-tracking analysis

The eye-tracking data was only minimally preprocessed. The data from each participant was zero-centered (using the overall mean over all trials), and data points with a Euclidean distance larger than 100 pixels (corresponding to 2.17° visual angle) from the zero-center were excluded from analysis (Fig. 1c and Fig. 5 show data before this exclusion). We analyzed the data in two epochs of interest, one time-locked to Stimulus 1 onset (from -500 ms until the onset of Test 1 at 5850 ms), and the other time-locked to Cue 2 onset (from -500 ms until the onset of Test 2 at 2850 ms). After artifact exclusion, on average 97.87% (SD = 1.50%, first epoch) and 95.14% (SD = 3.67%, second epoch) of the data remained for analysis.

#### Representational Similarity Analysis (RSA)

RSA of the gaze position data was performed separately for each participant using a single-trial approach. For each trial, we first obtained the trial-average for each of the 16 orientations while leaving out the current trial. We then computed at each time point the 16 Euclidean distances between the gaze position in the current trial and the trial-averages formed from the remaining data. This yielded a representational dissimilarity vector (RDV) of the distances between the (single-trial) gaze associated with the orientation in the current trial and the (trial-averaged) gaze associated with each of the 16 orientations (Fig. 2b). To examine orientation encoding, we computed at each time point and for each trial the Pearson correlation (rho) between the empirical RDV and the theoretical RDV predicted under a model of orientation encoding (see below) for the orientation on the current trial. When averaged over trials (and hence also across orientations), the procedure yields a leave-one-out cross-validated time-course of orientation encoding, similar to more conventional RSA approaches with trial-averages. However, the single-trial approach additionally retains the trial-by-trial variability in orientation encoding (Fig. 2b, *right* and Fig. 2e).

To examine orientation encoding within- and between objects (Fig. 3a), we used the same approach, but obtained the 16 trial averages separately for each of the 3 different objects in the participant’s stimulus set. This yielded 3 empirical RDVs per trial (one within- and two between-objects) that were independently correlated (Pearson’s rho) with the model RDV. The two between-objects correlations (rho’s) were then averaged.

#### Model geometries

Our basic orientation model was a perfect circle geometry (see Fig. 2a, left), where the model RDVs reflected the pairwise Euclidean distances between 16 evenly spaced points on the unit circle (see Fig. 2a, right; note that each line in the distance matrix corresponds to the model RDV for a given stimulus orientation; see Fig. 2b). The geometry of this model corresponds to our behavioral analysis model with 𝐵 = 0 (i.e., *M_circle_*, unbiased). To examine bias in the gaze patterns (Fig. 4), we used the Euclidean distance structures associated with our maximally biased models with 𝐵 = -1 (*M_attraction_*) and 𝐵 = 1 (*M_repulsion_*), respectively.

Comparing these two extreme models (which both have a square geometry) yields an estimate of the extent to which the gaze patterns were repulsively or attractively biased (see *Results*). Note that the distance structures expected under the three different models (𝐵 = 0, 𝐵 = 1, and 𝐵 = -1) correlate with each other (r = 0.77 and 0.34). We thus did not expect very large differences in their fit of the data and report the results with a more liberal statistical threshold (p_cluster_ < 0.05). In all cluster-based permutation tests (Maris & Oostenveld, 2007), we first identified clusters of consecutive samples that showed an effect with p_sample_ < 0.05 (uncorrected) and used the sum of t-values within a cluster as its test statistic. We then estimated the probability p_cluster_ that a cluster with a larger test statistic would emerge by chance, based on 20000 iterations where the individual subject effects were randomly sign-flipped. Unless otherwise specified, all reported statistical tests were two-sided.

#### Microsaccade detection

For complementary analysis of microsaccades (Fig. 5), we used a velocity-based detection algorithm established in previous work (De Vries et al., 2023; De Vries & Van Ede, 2023; Liu et al., 2023). In brief, the gaze position data were transformed into a velocity time course by calculating the Euclidean distances between consecutive samples, and smoothing it with a 7 ms Gaussian kernel. Saccade onsets and endpoints were inferred from when the gaze velocity exceeded a trial-specific threshold (5 times the median velocity within the trial) and when it returned to below threshold, with a minimum interval of 100 ms between successively detected saccades.

## Data and code availability

All data and code supporting this study are available at https://gin.g-node.org/lindedomingo/mpib_memoreye

## Author contributions

Juan Linde-Domingo: Conceptualization, Data Curation, Formal Analysis, Investigation, Project administration, Validation, Visualization, Writing – original draft, Writing – review & editing Bernhard Spitzer: Conceptualization, Funding acquisition, Methodology, Project administration, Resources, Supervision, Writing – original draft, Writing – review & editing

## Acknowledgments

We thank Ivan Padezhki, Clara Wicharz, Josefine Hebisch, Anna Faschinger, Gabriele Inciuraite and Antonia Anouk Bielefeldt for their help with data collection, and Jann Wäscher for participant recruitment. We also thank Josefine Hebisch for recording the auditory stimuli, Martin Rolfs for helpful comments and discussion, Ralph Hertwig for general support, and Thomas Graham for editorial assistance. This research was supported by a European Research Council (ERC) Consolidator Grant ERC-2020-COG-101000972 (BS) and by DFG grant SP 1510/7-1 (BS).

